# Presence of *Aedes aegypti* in a high-altitude area in Bolivia

**DOI:** 10.1101/2023.08.07.552199

**Authors:** César Ríos, Ninoska Rosas, Maria de los Ángeles Delgadillo-Iglesias, Maria Teresa Solis-Soto

**Author notes:** **Corresponding autor:**. Email address: CR NR MD.

## Abstract

*Aedes aegypti* is currently widespread in Bolivia. Although these mosquitoes commonly inhabit tropical and subtropical areas at low altitudes, recent studies found them up to 2,500 meters above sea levels (m.a.s.l.). The study’s objective was to identify the presence of the *Ae. aegypti* mosquito in urban areas with altitudes around 2,900 m.a.s.l. in Sucre, Bolivia. For this, ovitraps were installed at 12 points, and a morphological and taxonomic analysis was performed. The presence of mature and immature forms was confirmed in 3 of the 12 sampling points. Entomological indexes reported: House=13.64, Breteau= 27.27, dry containers= 43.55, containers with water =56.45.

Reproductive populations of *Ae. aegypti* was confirmed to more than 2,900 m.a.s.l. in an urban area with a high migratory flow area. It increases the risk for autochthonous cases of Dengue Disease in the short term. Prevention and control strategies must be intensified to reduce the transmission of mosquito-borne diseases.

## Introduction

Dengue is a viral infection caused by *Flaviviridae* which are transmitted to humans through the bite of infected mosquitoes. The main vectors that transmit the disease are *Aedes aegypti* mosquitoes and, to a lesser extent, *Ae. albopictus*[1]. In 2012, dengue was classified as the world’s most important mosquito-borne viral disease. It is the most prevalent arbovirosis in the last 50 years[2], and is considered endemic in more than 100 countries including Bolivia[1].

The global incidence of dengue has grown dramatically, with nearly half of the world’s population at risk. Although it is estimated that between 100 and 400 million infections occur yearly, more than 80% are generally mild and asymptomatic[1]. The determining factors for the evolution of the disease are not well known. Studies showed that children, secondary infection, ethnicity, and the presence of chronic diseases (diabetes mellitus, bronchial asthma, anemia) could be related to more severe disease cases[3]. It has been reported that infection with one of the serotypes could provide lifelong immunity against that serotype but only short-term immunity against other serotypes[4]. Likewise, sequential infection with different serotypes can trigger a more severe disease presentation[5].

*Aedes aegypty* is found mainly in tropical and subtropical areas. It has been reported that the climatic factor, especially temperature, directly influences the biology of the vector’s life cycle, favoring its reproduction and the incidence and prevalence of the disease. Temperatures higher than 31°C could accelerate its aging and mortality. On the other hand, temperatures lower than 21°C could result on the virus being more infectious due to the extended development and life duration. It seems that the daily variation in temperature is also a relevant factor in their life cycle, which has become more evident in the last decades[2, 6]. Studies have shown that dengue incidence increased by 2.6% one week after each 1ºC increase in maximum temperature and by 1.9% two weeks after each centimeter increase in precipitation[7]. Due to global warming and multiple regional and local socioeconomic factors, the distribution patterns of *Ae. aegypti* has changed, with an extensive distribution on all continents (except Antarctica)[8] and new altitudinal records reporting their presence in altitudes up to 2,550 m.a.s.l.[9]. Additionally, the rapid growth in urban populations can favor the proliferation of the vector, mainly due to inadequate management of solid waste, favoring reservoirs (e.g., stagnant water)[10, 11, 12].

The COVID-19 pandemic has caused unprecedented societal disruption, indirectly affecting the dynamics of infectious diseases[13], with significant consequences for at-risk populations[1]. Although the reduction in human mobility could explain the 44% decrease in the incidence of Dengue since March 2020 in some countries[13], mosquito control measures have also been interrupted during the pandemic, which could explain the rebound of cases currently, especially in the Americas[14, 15].

In Chuquisaca, Bolivia, the most vulnerable areas for vector proliferation are distributed along the Sancho (Río Chico), Pilcomayo, and Grande rivers, located along an extensive valley of intensive agricultural and recent tourist activity. This rural area presents a xeric-type bioclimatic zone, thermotropic ecological floor, with temperatures between 15.2 to 31.2 °C, annual accumulated precipitation of 948 mm^3^, and an altitude between 1,500 to 2,000 m.a.s.l.[16] There have been dengue outbreaks in several communities in recent years. On the other hand, the urban area of Sucre presents a higher altitude, colder temperatures, and less rain, without having reported the presence of the vector up to now. Their proximity of Rio Chico to Sucre’s urban area constitutes a vector migration risk. This situation, added to climate change and the consequences of control measures during the Covid-19 pandemic, made it necessary to carry out active vector surveillance measures in a larger area than the usual control. In that sense, this study aimed to identify the presence of the *Ae. aegypti* mosquito in urban areas with altitudes around 2,900 m.a.s.l. in Sucre as a potential risk of dengue disease transmission.

## Materials and Methods

### Setting and study design

A descriptive, cross-sectional study was carried out, capturing mosquito larvae between March 30 and April 20, 2022, in different parts of urban areas (District 2) in Sucre, Bolivia. Sucre is a city located in the central zone of Bolivia, in the inter-Andean valleys, with a population of 261,201 inhabitants[16]. It is situated at an altitude between 2.700 - 3000 m.a.s.l. with a sub-humid mountain climate with average minimum temperatures recorded in winter (July) of approximately 4.6 °C and average maximum temperatures of 24.3 °C in summer (November) and an average annual rainfall of 679 mm^3^[16].

### Sampling strategy

Sampling was performed considering the rainy months (epidemic period). It was carried out during the peak period of rainfall (29.03.2022 - 20.04.2022), reporting an average temperature of 14.6° C in this period.

Active entomological surveillance was performed using ovitraps at 12 strategic points in Sucre (Figure 1). Risk areas were identified according to REDILA protocol to determine the sampling point and the number of sites to install the ovitraps[17]. For field sampling, standardized field entomological techniques were used by the Vector-Transmitted Diseases Area of the Regional Health Service (ETV-SEDES Chuquisaca used to prevent and control mosquito-borne diseases). It consists of selecting the breeding sites within each strategic sampling point considering stagnant water and artificial environments (e.g. tires). The vector’s presence, density, and dispersion were reported at each sampling point.

**Figure 1.**
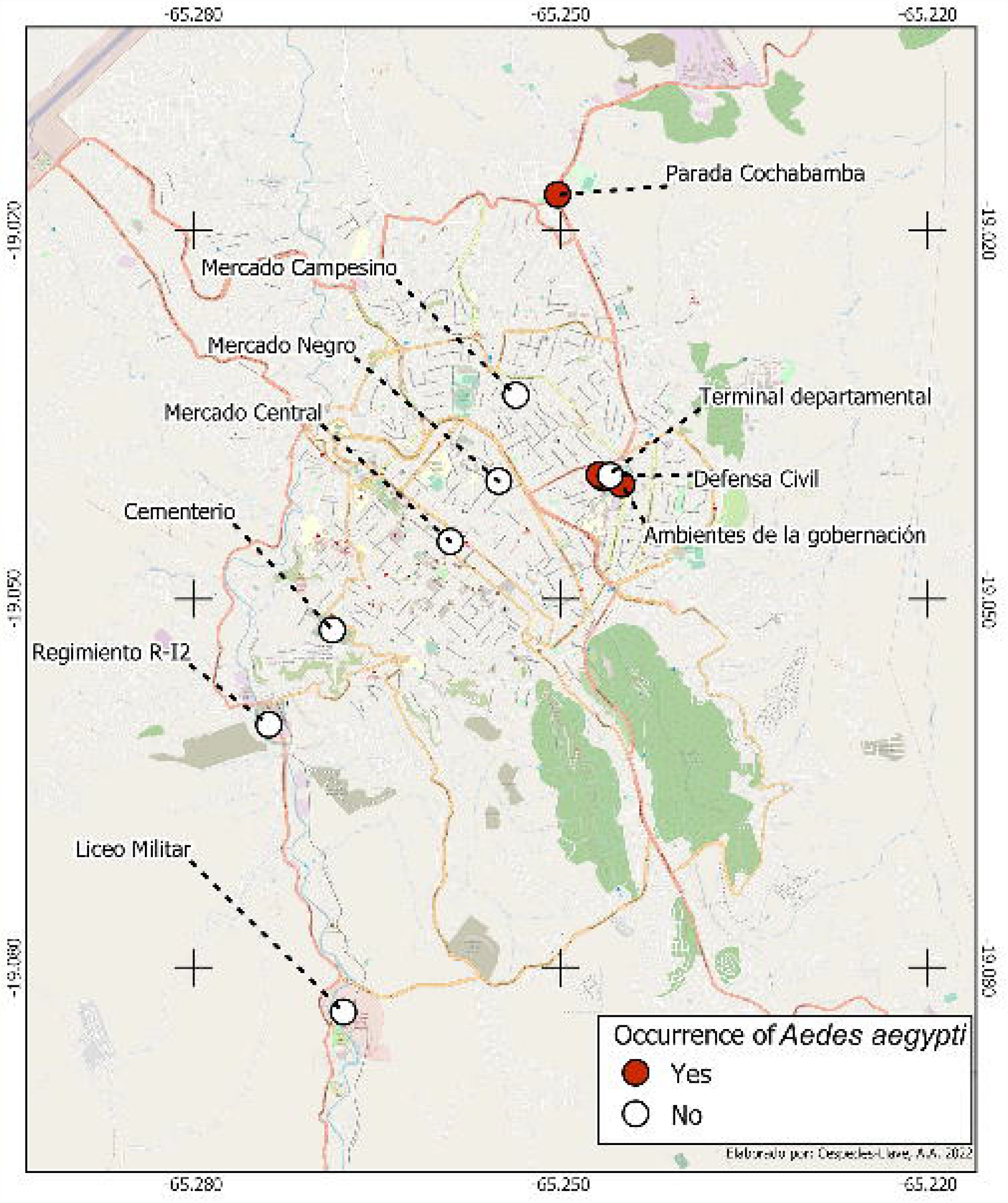
Sampling points (N=12), and the presence of ***Aedes aegypti*** in Sucre, Bolivia. (Terminal departamental and Ambientes de la gobernación, two sampling points each)

After the sampling stage, control actions were carried out at all sampling points (elimination of reservoirs, application of biological control with *Bacillus thuriengiensis*) and population education.

### Sample processing and analysis

The suspicious samples were processed according to National standardized protocols and sent to the entomology Laboratory of the National Center for Tropical Diseases (CENETROP) to confirm genus and species (morphological and taxonomic analysis). The larvae and the pupae stages were submerged in 70% alcohol, and the adult forms were stored in tubes. Entomological indices (House, Container, and Breateau) were computed as recommended by the Word Health Organization[19]. Taxonomic identification was carried out by using the taxonomic key of Rueda, 2004[18] to determine suspicious larvae, pupae, and adult stages of *Aedes aegypti*.

## Results

Three of the 12 sampling points (points 2, 4, 7) reported the presence of containers (tires with accumulated water), corresponding to 22 dwellings. Of these, 40 samples containing larvae were examined (Table 1). After the entomological recognition, mature (15 adults) and immature (five samples of larvae and five samples of pupae) forms suspicious of the mosquito *Ae. aegypti* were identified. A box with 15 adult culicids hatched in the Basic Entomology Unit, Sucre (UBE), sending mature and immature forms of the mosquito to the CENETROP for confirmation. The laboratory confirmed that 100% of the samples as *Ae. aegypti*. The entomological indices reported values higher than those reported for low transmission of the vector: (House index 13.64; Breteau index 27.27; Index of dry containers 43.55; and Index of containers with water 56.45).

**Table 1.** Entomological sampling for *Aedes aegypti*.

| N° | Altitud<br>(m.a.s.l.) | Place | Start date<br>(dd.mm) | End date<br>(dd.mm) | N° of Households | N° of containers |  | Positive collected samples |  | Samples analyzed | Positive samples |
| --- | --- | --- | --- | --- | --- | --- | --- | --- | --- | --- | --- |
|  |  |  |  |  |  | Dry | Wet | Dry | Wet |  |  |
| 1 | 2,798 | Cementerio General | 29.03 | 30.03 | 1 | 931 | 1132 | 0 | 0 | 5 | 0 |
| 2 | 2,837 | Terminal de buses departamental | 01.04 | 0.04 | 1 | 3 | 12 | 0 | 3 | 8 | 3 |
| 3 | 2,844 | Terminal de buses interprovincial | 01.04 | 0.04 | 1 | 4 | 8 | 0 | 0 | 0 | 0 |
| 4 | 2,834 | Ambientes de la gobernación | 05.04 | 05.04 | 1 | 13 | 201 | 0 | 2 | 9 | 2 |
| 5 | 2,833 | Defensa civil | 06.04 | 06.04 | 1 | 1 | 1 | 0 | 0 | 0 | 0 |
| 6 | 2,833 | Gobernación (Garaje) | 06.04 | 06.04 | 1 | 10 | 18 | 0 | 0 | 2 | 0 |
| 7 | 2,933 | Salida a Cochabamba | 13.04 | 13.04 | 11 | 215 | 157 | 0 | 1 | 13 | 1 |
| 8 | 2,650 | Liceo militar | 19.04 | 19.04 | 1 | 32 | 11 | 0 | 0 | 1 | 0 |
| 9 | 2,694 | Regimiento R-12 | 19.04 | 19.04 | 1 | 5 | 12 | 0 | 0 | 1 | 0 |
| 10 | 2,788 | Mercado Central | 20.04 | 20.04 | 1 | 26 | 47 | 0 | 0 | 0 | 0 |
| 11 | 2,805 | Mercado Negro | 20.04 | 20.04 | 1 | 1 | 5 | 0 | 0 | 0 | 0 |
| 12 | 2,803 | Mercado Campesino | 20.04 | 20.04 | 1 | 2 | 7 | 0 | 0 | 1 | 0 |
| **Total** |  |  |  |  | 22 | 1243 | 1611 | 0 | 6 | 40 | 6 |

Georeferencing of positive points:
Point 2 (1): Latitude -19.040118; Longitude -65.246530
Point 2 (2): Latitude -19.039900; Longitude -65.246880
Point 4: Latitude -19.040650; Longitude -65.245036
Point 7: Latitude -19.017078; Longitude -65.250217

In the morphological analysis, lateral spines were found in five larvae at the base of the group of bristles of the mesothorax, as well as scales and detail in the eighth segment (Picture 1). In the taxonomic analysis of the adult forms, six females and nine males were identified, all with the presence of the characteristic lyre *Ae. aegypti* in the thorax (Picture 2).

## Discussion

We present the results of the entomological surveillance of the transmitting vector of Dengue disease in urban areas of Sucre in Bolivia. The results confirm the presence of this vector at an altitude that has not been previously reported (greater than 2,900 m.a.s.l.).

Previous studies have reported some adaptations of the vector under unusual conditions. *Ae. aegypti* mosquito was identified in some high-altitude areas, mainly in Asia and Latin American countries. Immature forms of the mosquito were reported in eastern Nepal (up to 2,000 m.a.s.l.)[20] High Mountain region of Nepal, the Central Himalayas (between 85 to 2,100 m.a.s.l.)[21], Puebla city in Mexico (2130 m.a.s.l.)[22], Mexico City (2,250) and Colombia (2,302m.a.s.l.)[23]. Our results are also consistent with a previous study in the city of Cochabamba, Bolivia, where the presence of the vector was reported up to 2,550 m.a.s.l. in a period of high rainfall, favoring the generation of common and uncommon artificial breeding sites[9]. Identifying this vector at 2,933 m.a.s.l. corroborates their rapid adaptation. It could become a risk factor for autochthonous transmission in cities located even at high altitudes (almost 3,000 m.a.s.l.) due to global warming in the short term.

Although the entomological indices have shown weak associations between vector indices and dengue transmission[24], our results show higher values, more than double, than those reported. It suggests implementing systematic routine control activities[25].

Likewise, this distribution change could be partly explained as an indirect consequence of the Covid-19 pandemic. Disruption to routine vector control programs (e.g. spraying)[26] and changes in mosquito-human interaction have been reported in many countries, which may have favored the migration and proliferation of the vector into new geographical areas[27].

On the other hand, in Chuquisaca, Bolivia, as in other countries, it has not been possible to fully implement the integrated management strategies for arbovirosis (EGI-ARBOVIROSIS), suggested by the Pan American Health Organization for cities without mosquitos [28]. It increases the communities’ vulnerability even more when the dynamics of the vector are not being considered in work with the communities, and they are unfamiliar with the strategies to control the transmission of dengue, zika, and chikungunya.

The study has some limitations. Due to limited resources, it could not cover more sampling points nor repeat it for more days. It limited the possibility of capturing more mosquitoes in its different phases. Although the study could not determine the presence of arboviruses, the presence of this mosquito in crowded and highly mobile areas, such as the bus terminal and public markets, makes us reflect on the potential risk of transmission in humans.

The presence of *Ae. aegypti* to more than 2,933 m.a.s.l. in an urban area with a high migratory flow from and to endemic regions of Dengue disease recognizes an increased risk for autochthonous cases or mosquito adaptation in the short term. Due to the effects of climate change, having dry seasons, and areas lacking water supply networks, some common practices (e.g., storing containers with rainwater) increase the vector’s proliferation risks. In this sense, prevention and control strategies must be intensified, mainly in endemic areas, as well as surveillance and early warning systems to reduce the transmission of mosquito-borne diseases such as dengue.

## Acknowledgments

We want to thank the entomology technicians of the Departmental Health Service (SEDES), Chuquisaca René Rentería, Santos Chonta, and Carlos Plaza.

## Availability of data and materials

The data are available on request.

## Authors’ contributions

CR, NR, MTSS They made substantial contributions to the analysis, interpretation of the data, and writing the draft of the manuscript. MD participated in the interpretation of the data. All authors read and approved the final version of the manuscript.

## Ethics approval and consent to participate

Not applicable.

## Consent for publication

Not applicable.

## Competing of Interest

The authors declare no competing of interest.

**Picture 1.**
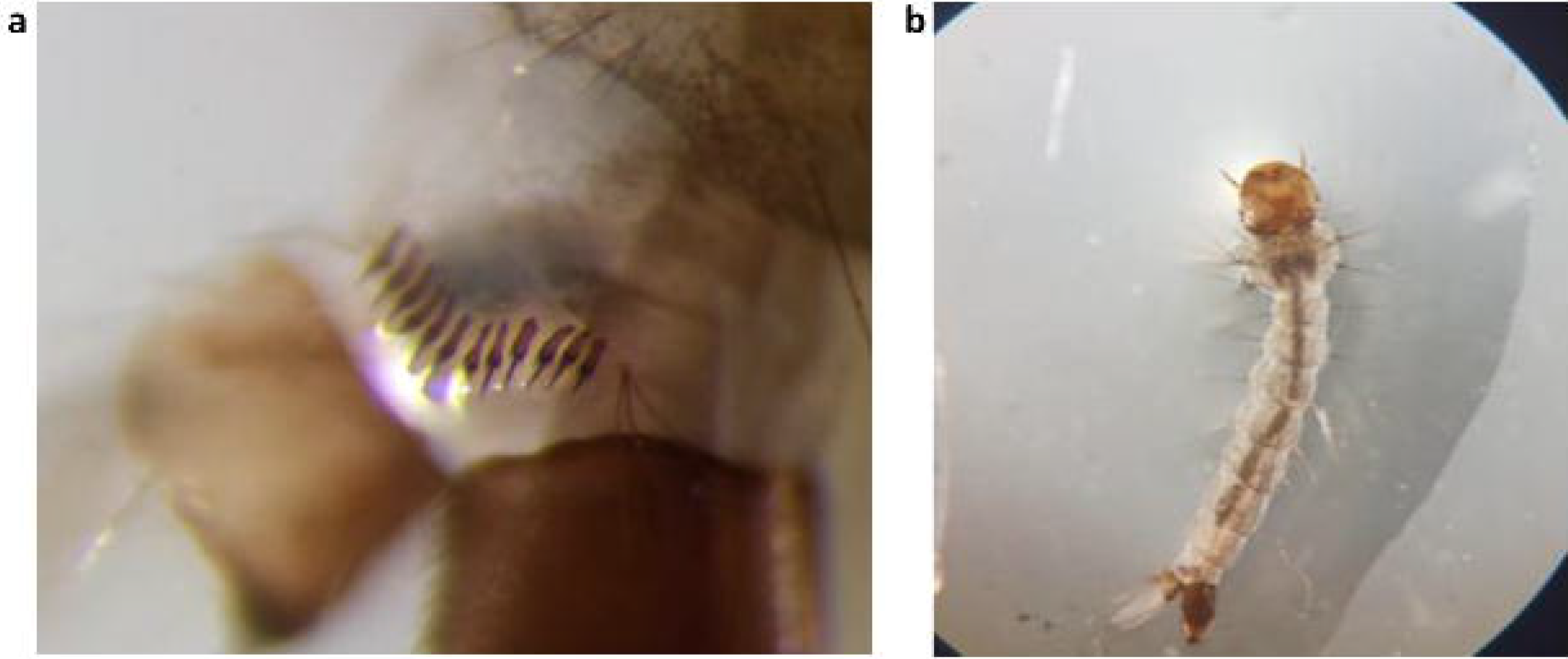
*Aedes aegypti* larvae. In a you can see the scales of the eighth second of the larva. b. Lateral spines are seen at the base of the metathorax bristle cluster.

**Picture 2.**
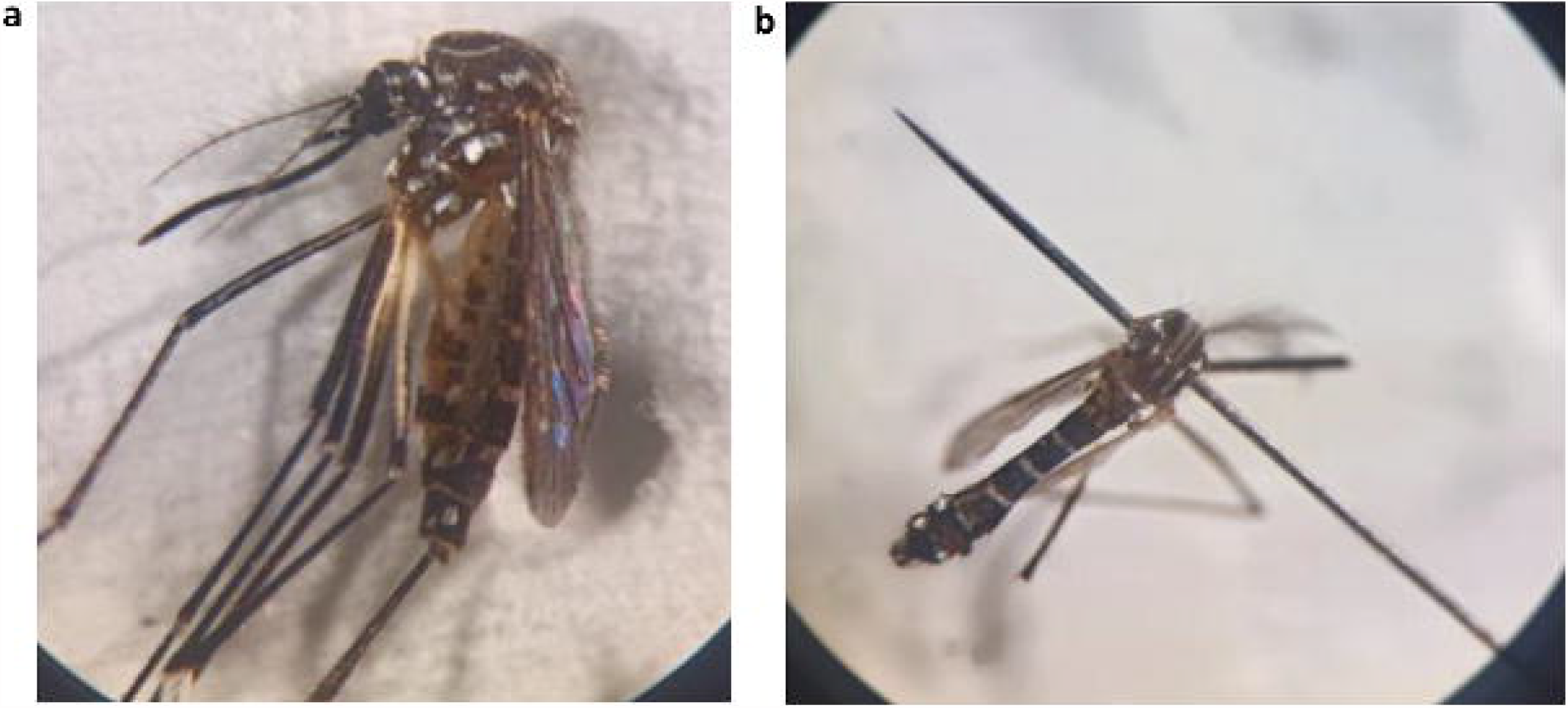
*Aedes aegypti* adults. a. *Aedes aegypti* female with side view. b. Top view of male *Aedes aegypti*

